# Climate and urbanization structure West Nile virus vector communities: Tools for surveillance under environmental change

**DOI:** 10.64898/2026.07.24.738033

**Authors:** Benjamin P. Gregory, Arielle Arsenault-Benoit, Patrick Irwin, Keith J. Price, Bryn J. Witmier, Matthew C. Fitzpatrick, Megan L. Fritz

## Abstract

West Nile virus (WNv) in eastern North America is transmitted primarily by *Culex pipiens* and *Culex restuans*, two cryptic vector species whose relative abundance shifts seasonally and across urbanization gradients - gradients that are themselves now shifting under continuing climate change and urban expansion. Because the two species are difficult to separate morphologically and are typically pooled in routine surveillance, this turnover is seldom tracked directly, and agencies lack species-resolved tools to anticipate when, where, and how vector communities will reorganize. Yet the timing of this turnover has been linked to the seasonal timing and intensity of human WNv cases, making it a potentially forecastable correlate of risk. We molecularly identified 9,789 *Culex* collected over six years, of which 5,808 came from a designed multi-year study in the Baltimore-Washington metropolitan region, and the remainder from Philadelphia and Chicago for cross-regional comparison. We combined negative-binomial generalized linear mixed models of species-specific abundance in Baltimore-Washington with Gradient Forest models of compositional turnover across all three regions. The two species diverged along both the urbanization and thermal gradients: *Cx. pipiens* abundance increased with impervious surface and with temperature, whereas *Cx. restuans* declined along both. Consequently, highly urbanized areas remained *Cx. pipiens*–rich across the season, whereas suburban, low-to-moderate-development landscapes exhibited the largest seasonal shifts in community composition. Accumulated degree-days (ADD) and weekly mean temperature were the most consistent drivers of turnover across regions, with the shift from *Cx. restuans* to *Cx. pipiens* fastest between 594 and 610 ADD. These patterns translate into three operational tools for WNv management: a climate-based degree-day window that lets agencies forecast the community shift from temperature data before it is detectable in trap composition, identification of suburban landscapes as priority targets for intensified surveillance and early intervention, and species-resolved environmental responses that help anticipate how community composition may shift within these regions as local climate and land-use conditions change. Together they offer a path from reactive surveillance to anticipatory, spatially targeted, and forward-looking management of WNv risk under environmental change.

**Open Research Statement:** All datasets and R scripts used to generate the results and figures in this study will be made publicly available on DRYAD (DOI pending) upon acceptance of the manuscript. Climate data are publicly available from the PRISM Climate Group (PRISM Group, Oregon State University). Landscape predictor datasets are publicly available from the sources cited in Methods (Dewitz 2023; WorldPop 2018; U.S. Census Bureau 2020; Zell & Sanford 2020; Gesch et al., 2018; Didan 2015).

## Introduction

West Nile virus (WNv) is the most widespread mosquito-borne virus in North America, maintained in an enzootic cycle between birds and *Culex* mosquitoes and amplified across much of the continental United States each summer (Andreadis, 2012; Kilpatrick, 2011). In the Northeastern United States, two cryptic, co-occurring species - *Culex pipiens* and *Culex restuans* - are its principal vectors (Andreadis, 2012; Rochlin et al., 2019). The two are difficult to separate as adults and are routinely pooled in surveillance programs, where they are reported together as “*Cx. pipiens/restuans*” and summarized as a single vector index alongside infection rates to assess seasonal WNv risk (CDC, 2024; Ebel et al., 2005; Harrington & Poulson, 2008; Johnson et al., 2015). But the two species differ in phenology, thermal physiology, and habitat associations, and the climatic and landscape gradients that shape their relative abundance are themselves now shifting under continuing warming and urban expansion (Ainsworth, 2023; Erazo et al., 2024; Gorris et al., 2021; Mordecai et al., 2019). This distinction is epidemiologically consequential: the timing of the seasonal shift from one species to the other has been associated with the timing and intensity of human WNv cases (Lampman et al., 2006; Tokarz & Smith, 2020). A pooled index cannot register how the two species are responding differently to climatic and land use changes, leaving surveillance programs without the species-resolved information they need to anticipate when and where vector communities will reorganize. Resolving how each species responds to temperature and urbanization - separately, and across more than one region - is therefore both an ecological and an operational question of growing relevance under environmental change.

The phenological contrast between the two species has a physiological basis: *Cx. restuans* develops more rapidly at lower temperatures and suffers higher mortality above ∼25 °C, while *Cx. pipiens* develops faster and survives better under warmer conditions (Buth et al., 1990; Ciota et al., 2014; Madder et al., 1983). These thermal differences manifest as a seasonal succession in which *Cx. restuans* abundance peaks earlier under cooler conditions and *Cx. pipiens* reaches higher abundance later in the summer as temperatures rise (Andreadis et al., 2001; Arsenault-Benoit & Fritz, 2023; Johnson et al., 2015; Lee & Rowley, 2000), placing *Cx. pipiens* at peak abundance during the late-summer window when WNv has amplified in avian reservoirs and human cases begin to appear (Kunkel et al., 2006; Lampman et al., 2006; Tokarz & Smith, 2020). By contrast, the two species are broadly similar in their host use and laboratory vector competence. Both feed predominantly on birds, take blood meals from humans at similar low frequencies, and are competent WNv vectors in the laboratory (Ebel et al., 2005; Rochlin et al., 2019). Yet they contribute differently to field WNv transmission across the season: *Cx. restuans* is most abundant and frequently WNv-infected earlier in the summer, consistent with a role in early-season enzootic amplification, whereas *Cx. pipiens* predominates later, during the period when human cases occur (Burgunder et al., 2026; Kunkel et al., 2006; Lampman et al., 2006; Tokarz & Smith, 2020).

How environmental gradients structure these communities is reasonably well understood for climate but less so for landscape, and rarely for both together. Temperature and accumulated thermal units are reliable predictors of *Culex* seasonal dynamics and of the transition between species (Ciota et al., 2014; Kunkel et al., 2006; Lebl et al., 2013; Yu et al., 2018). Landscape structure also shapes their distributions, though the evidence is mixed: several studies report habitat partitioning, with *Cx. pipiens* favoring highly urbanized, impervious environments (Arsenault-Benoit & Fritz, 2023; Bondo et al., 2023; Gorris et al., 2021; Johnson et al., 2015) and *Cx. restuans* associated with more vegetated or less developed habitats (Arsenault-Benoit & Fritz, 2023; Gorris et al., 2021; Johnson et al., 2015), whereas others find broadly generalist habitat use with little differentiation (Ebel et al., 2005; Gardner et al., 2012; Irwin et al., 2008). Critically, most of this work is confined to a single metropolitan area or a single season - for example, five suburban New Jersey sites sampled in one year (Johnson et al., 2015), or a single urbanization gradient in metropolitan Washington (Arsenault-Benoit & Fritz, 2023) - making it difficult to separate climatic from landscape effects or to judge how far any pattern generalizes across regions. This matters because WNv risk depends jointly on total vector abundance, which sets the magnitude of transmission potential (Garrett-Jones, 1964), and on species composition, which determines which vector predominates as the season advances (Burgunder et al., 2026; Lampman et al., 2006; Tokarz & Smith, 2020). Resolving either across regions, and especially their interaction, requires data and methods that few existing studies can offer.

The gradients structuring these communities are shifting. West Nile virus has expanded northward in Europe over the past decade, driven in part by warming temperatures that lengthen vector-activity seasons and enlarge the area climatically suitable for virus circulation (Ainsworth, 2023; Erazo et al., 2024). In North America, the ranges of key *Culex* vectors are projected to follow (Gorris et al., 2021; Samy et al., 2016) and continuing urban and suburban development creates habitat for human-commensal *Culex* lineages in regions where they were previously absent (Haba et al., 2025; Mutebi & Savage, 2009). Because *Cx. pipiens* and *Cx. restuans* differ in thermal physiology and habitat associations, the same temperature and land-use changes that broaden vector distributions will also alter the relative balance between the two species, and with it, the seasonal phenology of the community. A pooled index sums two responses likely to move in opposite directions and so cannot register this rebalancing. Anticipating how vector communities will reorganize and where surveillance and control effort should follow therefore requires environmental responses fit to each species individually, characterized across enough of the climatic and landscape gradient to generalize beyond any one region (Mordecai et al., 2019).

Here we integrate a multi-year, designed sampling effort in one focal region with complementary samples from two additional regions to characterize how climate and urbanization jointly structure *Culex* vector communities, and to translate that structure into operational guidance for surveillance under continuing environmental change. We molecularly identified *Culex* collected over six years across the Baltimore-Washington metropolitan region - where balanced spatial and temporal sampling supported both compositional and species-specific abundance modeling - together with samples from Philadelphia and Chicago, drawn from existing operational and research collections that allowed us to test whether the compositional turnover patterns observed in Baltimore-Washington were consistent with those observed in independent regions sampled under different designs. We analyzed the data in two ways. First, in the Baltimore-Washington region, where sampling was spatially and temporally balanced, we used generalized linear mixed models to evaluate how temperature and urbanization shape the abundance of each species and of the community as a whole. Second, we used Gradient Forest, a machine-learning approach that quantifies nonlinear compositional turnover along environmental gradients and identifies thresholds where composition changes most rapidly (Ellis et al., 2012; Fitzpatrick & Keller, 2015; Pitcher et al., 2011), to locate the thermal point - expressed in accumulated degree-days (Gu & Novak, 2006; Vinogradova, 2000) - at which the community shifts from *Cx. restuans* to *Cx. pipiens* and to test whether that threshold is consistent across regions. Within Baltimore-Washington, we used the same compositional approach, combined with a beta-binomial model of species composition, to map predicted relative abundance of the two species across the landscape and to identify the landscape conditions across which composition shifts most rapidly. Together, these analyses give vector-control programs three things: a transferable, climate-based threshold for anticipating when the seasonal shift will occur, a spatially explicit description of where the shift is most pronounced on the landscape, and species-resolved environmental responses that help anticipate how community composition may shift within the regions studied as local climate and land-use conditions change.

## Methods

### Mosquito collection and identification

Mosquitoes were collected across three eastern metropolitan regions, with substantial differences in study design. Baltimore-Washington was sampled under a multi-year designed study optimized for spatial and temporal balance and supported both compositional and abundance analyses. The Philadelphia and Chicago datasets were drawn from existing surveillance and research collections that were not designed for this comparison. They supported compositional analyses across regions but did not yield the per-trap-night total catch data required for abundance modeling. Details of each locality follow (Fig. 1, DataS1).

**Figure 1.**
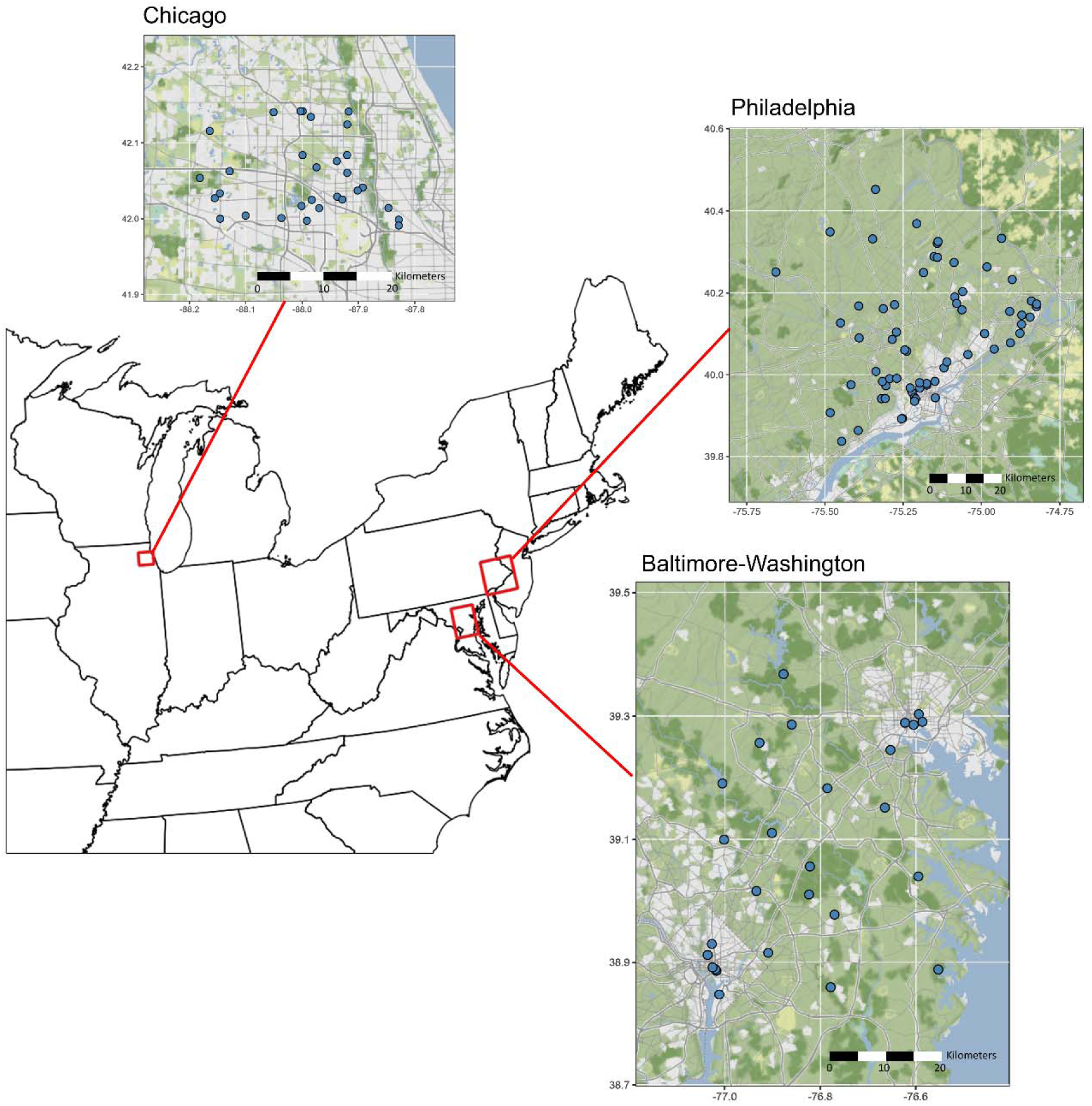
Collection sites in three metropolitan localities in the eastern United States: Northwest Chicago, Illinois; Philadelphia, Pennsylvania; and the Baltimore-Washington Metropolitan Region (BW).

### Baltimore-Washington (BW)

Fifteen collection sites were established in 2019, all within approximately 50 km of downtown Washington, District of Columbia. From 2019 through 2022, sites were visited every two weeks in a randomized order from June 1^st^ through September 30^th^, with the exception of 2021, when sampling started on April 1^st^. In 2023, three sites were dropped, and twelve new sites were added, bringing the total number of sites to 24 for 2023 and 2024 (but 27 for the whole dataset). In 2023 and 2024, sites were visited once per month in a randomized order between June 1 and October 1. The size of the total sampling area was 2,662 km^2^.

### Chicago

In Chicago, nine sites regularly surveyed by the Northwest Mosquito Abatement District (NWMAD) were selected and established for this study. Specimens were collected biweekly on eight occasions per year from June to October in 2020 and 2021. As part of a separate study (Burgunder et al., 2026), 21 additional sites in the NWMAD were sampled for three consecutive collection dates in July 2021, four consecutive dates in August 2021, and five consecutive dates in both July and August of 2022. All sites were approximately 50 km northwest of downtown Chicago, IL, and encompassed a 511 km^2^ region in northwestern Cook County, IL.

### Philadelphia

In Philadelphia, specimens were collected as part of the vector surveillance and management activities of the Pennsylvania Department of Environmental Protection Division of Vector Management (PA DEP). In 2019, mosquitoes collected in Bucks, Montgomery, Delaware, and Philadelphia Counties, PA, were retained on a weekly basis from May 23, 2019 to August 20, 2019. The number of mosquitoes within each collection varied, with a minimum of ten per trap. In 2020 and 2021, up to 10 samples were retained per county per month from May through September in Philadelphia and Bucks counties. A total of 65 sites were distributed across a 4,885 km^2^ region, covering much of southeastern Pennsylvania.

All specimens were collected via CDC gravid traps (John W. Hock Company, Gainesville, FL, USA, Model 1712) set overnight with an alfalfa-infused oviposition attractant. Because trap type, attractant, and overnight deployment were held constant across all sites and dates, gravid trap catch can support inference about how abundance *varies* across spatial and climatic gradients. Gravid traps are also well suited to *Culex* surveillance (DiMenna et al., 2006; Karki et al., 2016; Williams & Gingrich, 2007), and our two focal species, *Cx. pipiens* and *Cx. restuans*, are equally attracted to the alfalfa infusion (Lampman & Novak, 1996).

Dates and GPS coordinates for each collection can be found in DataS1. Adult mosquitoes suspected to be *Cx. pipiens* or *Cx. restuans* were separated from non-*Culex* species morphologically and stored at −80°C until molecular identification at the University of Maryland, College Park. Because the two species are difficult to distinguish morphologically in their adult stage (Harrington & Poulson, 2008), PCR-based species-diagnostic markers have been developed (Crabtree et al., 1995; Smith & Fonseca, 2004). Our specimens were molecularly identified as described in Arsenault-Benoit and Fritz (2023); a full description of these methods can be found in Appendix S1. Abundance modeling requires the total overnight trap catch, which was retained only for Baltimore-Washington; in Philadelphia and Chicago, only a subsample of each catch was retained for genotyping, yielding species proportions but not total abundance. We therefore used the Philadelphia and Chicago data only for compositional (Gradient Forest) analyses and restricted abundance modeling to Baltimore-Washington.

### Quantification of landscape predictors

Landscape predictors were chosen due to their influence on mosquito distribution as identified by previous research (Diuk-Wasser et al., 2006; Gorris et al., 2021). The predictors used were percent impervious surface and percent tree canopy cover at resolutions of 30m (Dewitz, 2023), as well as log-transformed population density and housing unit density at a resolution of 30 arcseconds (WorldPop, 2018; U.S. Census Bureau, 2020), water table depth at a resolution of 250m (Zell & Sanford, 2020), elevation at a resolution of 30m (Gesch et al., 2018), and Normalized Difference Vegetation Index (NDVI) at a resolution of 250m (Didan, 2015). The value for each predictor was measured within a buffer zone around each site with a 1 km diameter using the R package terra (Hijmans, 2020). This buffer size was selected because the mean mark-recapture distance for this genus is approximately 1 km (Ciota et al., 2012; Hamer et al., 2014).

### Quantification of climatic predictors

Daily climate data were retrieved from the PRISM database (PRISM Group, Oregon State University) for each trap-night’s coordinates and date at a resolution of 800m. Climate data included daily precipitation (mm), daily mean temperature (°C), and daily maximum temperature (°C). Climate predictors were aggregated across weekly time lags to determine the mean for each predictor one to seven days and eight to 14 days preceding the collection date. Climatic conditions over these lags have been cited as reliable predictors of *Cx. pipiens*-*restuans* populations and WNv cases (Lebl et al., 2013). Finally, accumulated degree-days (ADD) was calculated for each trap-night using daily mean temperature data, as in Gu and Novak (2006), using the following formula:

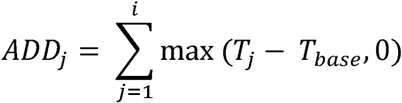

where T*_base_* is the baseline development temperature for *Culex pipiens* (10°C) (Vinogradova, 2000) and T*_j_* is the mean temperature on day *j* We used a single, *Cx. pipiens*–based threshold for two reasons. First, Gradient Forest characterizes turnover along a single accumulated–degree-day gradient, so a common base temperature is required to place both species on one axis. Second, the resulting degree-day window indexes the timing of the seasonal transition to higher *Cx. pipiens* abundance - the event of operational interest for surveillance - making the *Cx. pipiens* developmental base the appropriate referent. *Culex restuans* has a lower developmental threshold (embryonic ≈ 4.4 °C, larval ≈ 9.4 °C; Madder et al., 1983); recomputing degree-days on a *Cx. restuans* base would shift the numerical value of the threshold but not its interpretation or operational use.

### Abundance modeling

We modeled *Culex* trap-night abundance in BW using negative binomial generalized linear mixed models (GLMMs) with a log link. We fit site, year, and trap-night as random intercepts to account for repeated sampling across space and time and shared variation among traps run on the same night. Variance components, model fit (marginal and conditional *R*²; trigamma approximation, Nakagawa et al., 2017), negative binomial dispersion, and grouping-factor sample sizes for both models are reported in Table S1. R code for these diagnostics was developed with assistance from Claude Opus 4.7 (Anthropic) in June 2026 and reviewed and validated by the authors. The species-factor model was refit under alternative optimizers (Nelder–Mead, nlminbwrap) to confirm robustness of fixed-effect estimates. The negative binomial distribution accommodated overdispersion, and continuous predictors were centered and scaled (mean = 0, SD = 1). For both the overall and species-specific analyses, we selected predictors by two-stage forward selection on AIC. We first fit every single-predictor model and retained the lowest-AIC predictor, then anchored on it and fit every two-predictor model - including predictor interactions - retaining the lowest-AIC combination. Within each screen, all models were fit to a common set of complete-case observations so that AIC values were comparable. Candidate predictors were the landscape and climatic variables described above: percent impervious surface, percent tree canopy, NDVI, elevation, population density, housing density, water table depth, weekly and lagged-weekly mean and maximum temperature, weekly and lagged-weekly precipitation, accumulated degree-days, and day of year. Full screening results are reported in Tables S2-S5. The distributions of these landscape predictors across sites within each region are shown in Figure S1.

### Combined Culex abundanc

We modeled total abundance (*Cx. pipiens* + *Cx. restuans* per trap-night) against each scaled predictor. Single-predictor models contained one predictor; the best-supported was elevation. Two-predictor models added each remaining predictor and its interaction with elevation; the best-supported added percent impervious surface. The final overall model therefore comprised elevation, percent impervious surface, and their interaction.

### Species-specific abundance

To test whether species responded differently to environmental gradients, we fit a single GLMM with one observation per species per trap-night (two observations per trap-night), with species (a two-level factor) interacted with each environmental predictor. This is one model with species-by-predictor interactions, not separate per-species models. For trap-nights with more than 30 *Culex*, species-specific counts were estimated by multiplying the total catch by the proportion of each species among a random subsample of 30 genotyped individuals. Single-predictor models included species and its interaction with one predictor; the best-supported predictor was percent impervious surface. Two-predictor models included species interactions with both predictors (all two- and three-way terms); the best-supported added weekly mean temperature. The impervious-surface × temperature and three-way interactions were non-significant and were removed, leaving species-specific effects of impervious surface and temperature.

We then added a species-specific elevation effect directly, rather than by extending the stepwise screen. This was a single, directed addition because elevation was the best-supported predictor of overall abundance (see Appendix S1 for full rationale). We confirmed that elevation was not strongly collinear with the predictors already in the model (strongest pairwise *r* = −0.50, elevation–impervious; elevation–temperature *r* = −0.13; variance inflation factors ≤ 1.4; Table S6), and that the species-specific elevation slopes were essentially unchanged whether or not temperature was included, indicating an elevation effect independent of temperature. Adding species-specific elevation improved fit over the impervious-surface-plus-temperature model (ΔAIC = 11.6; likelihood-ratio test χ² = 15.6, *p* < 0.001; Table S7), so we report the elevation-inclusive model as the species-factor model (Table 1). Models were fit with the bobyqa optimizer (Bates et al., 2015).

**Table 1.** Summary of negative binomial generalized linear mixed models (GLMMs) examining environmental predictors of overall *Culex* abundance and species-specific abundance in the Baltimore-Washington region. Models included random intercepts for site, year, and trap night. β represents fixed-effect coefficients on the log scale, SE is the standard error, and *z* and *p*-values correspond to Wald tests of fixed effects. Continuous predictors were centered and scaled prior to analysis. IRR (incidence rate ratio) is the exponentiated coefficient (expβ) and represents the multiplicative change in expected abundance associated with a one standard deviation increase in the predictor (or, for categorical predictors, relative to the reference category), holding other predictors constant.

| Model | Predictor | $\beta$ | SE | $z$ | p-value | IRR<br>($\exp\beta$) |
| --- | --- | --- | --- | --- | --- | --- |
| ***Culex* Overall** | **Intercept** | 2.091 | 0.250 | 11.580 | < **0.001** | 8.09 |
|  | **Elevation** | 0.166 | 0.101 | 1.639 | 0.101 | 1.18 |
|  | **Percent Impervious Surface** | -0.139 | 0.098 | -1.417 | 0.157 | 0.87 |
|  | **Elevation x Percent Impervious Surface** | -0.422 | 0.121 | -3.486 | < **0.001** | 0.66 |
| **Species-Specific** | **Intercept** | 2.713 | 0.262 | 10.37 | < **0.001** | 15.07 |
|  | **Species (*Cx. restuans*)** | -1.236 | 0.076 | -16.30 | < **0.001** | 0.29 |
|  | **Elevation** | 0.484 | 0.120 | 4.05 | < **0.001** | 1.62 |
|  | **Percent Impervious Surface** | 0.365 | 0.114 | 3.20 | 0.001 | 1.44 |
|  | **Weekly Mean Temperature** | 0.326 | 0.069 | 4.72 | < **0.001** | 1.39 |
|  | **Species x Elevation** | -0.240 | 0.085 | -2.83 | **0.005** | 0.79 |
|  | **Species x Percent Impervious Surface** | -1.312 | 0.091 | -14.47 | < **0.001** | 0.27 |
|  | **Species x Weekly Mean Temperature** | -0.798 | 0.076 | -10.50 | < **0.001** | 0.45 |

### Gradient forest modeling of species turnover in Baltimore-Washington

We used GF, a machine-learning approach designed to quantify the environmental drivers of community composition (Ellis et al., 2012; Pitcher et al., 2011), to model compositional turnover of *Cx. restuans* and *Cx. pipiens* across environmental gradients. We chose GF because it flexibly models nonlinear, interacting effects of multiple predictors on community composition and produces continuous cumulative turnover functions that describe the magnitude and rate of species compositional turnover along each predictor (Ellis et al., 2012; Pitcher et al., 2011).

The model was fit using 1,000 regression trees per species (*Cx. pipiens* and *Cx. restuans*) with the R package gradientForest (Ellis et al., 2012). A correlation threshold of 0.5 was set to avoid overestimation of importance metrics for correlated predictors (Strobl et al., 2008). Two predictors were randomly sampled as candidates at each split, and samples were split into training and testing sets at proportions of 0.63 and 0.37, respectively. The model was fit to trap-night–level proportion of *Cx. pipiens* (and complementary proportion *Cx. restuans*), using the same set of variables described above in the GLMMs: percent impervious surface, percent tree canopy, NDVI, elevation, population density, housing density, water table depth, weekly and lagged-weekly mean and maximum temperature, weekly and lagged-weekly precipitation, accumulated degree-days, and day of year as the predictors. R^2^-weighted importance values determined which environmental predictors were the most important drivers of species turnover. We assessed where along each predictor gradient the rate of turnover between species occurred most rapidly (potential ecological filters or thresholds) by calculating the maximum of the ratio of the density of splits to the density of the input data (Ellis et al., 2012, Fitzpatrick and Keller, 2015); R code for this extraction was developed with assistance from GPT-4 (OpenAI) in June 2025 and was reviewed and validated by the authors.

### Spatial predictions in Baltimore-Washington

Because of the spatial extent and duration of sampling in BW, we made spatial predictions of *Cx. pipiens* relative abundance across two phenological windows in this region. First, we split BW trap-night data into early (1,078 mosquitoes from 97 trap nights) and late (4,730 mosquitoes from 481 trap nights) seasonal windows based on the accumulated degree-day threshold (594 accumulated degree-days) identified by the Baltimore-Washington Gradient Forest (GF) model, described above. For each season-site combination, we calculated proportion of *Cx. pipiens* as the sum of mosquitoes genotyped as *Cx. pipiens* divided by total mosquitoes genotyped as either *Cx. pipiens* or *Cx. restuans* across all sampling timepoints. We then trained a separate GF model for each seasonal window, using the corresponding site-wise summaries of proportion of *Cx. pipiens* (and complementary proportion *Cx. restuans*) as the responses; and the landscape predictors described above (elevation, housing density, NDVI, population density, percent impervious surface, percent tree canopy, and water table depth). The resulting GF turnover functions were used to transform environmental predictors into biological importance values following Fitzpatrick and Keller (2015). We then performed a Principal Components Analysis (PCA) on the transformed predictors to generate uncorrelated spatial predictors. Because the PCA inputs were biological importance values rather than raw predictor units, PC1 is a composite axis on a biological-importance scale, weighted toward whichever predictors contributed most strongly to compositional turnover in the underlying GF model (Table S8); the variance it explains is variance in the transformed predictor space, not variance in the species-composition response.

To model species composition, we fit a generalized linear model with a beta-binomial error distribution and logit link, implemented in glmmTMB (Brooks et al., 2017), relating the number of *Cx. pipiens* individuals (out of the total number of genotyped *Culex*) to the first principal component (PC1) of the PCA. This model took the form:

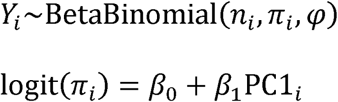

Where *y_i_* represents the number of *Cx. pipiens* individuals identified among *n_i_* genotyped *Culex* mosquitoes at site-season *i,*and *π_i_* represents the probability that a sampled mosquito belongs to *Cx. pipiens*. The parameter φ describes extra-binomial dispersion. The beta-binomial distribution was used to accommodate overdispersion while preserving the binomial count structure. The fitted model was used to predict the expected proportion of *Cx. pipiens* across the study extent for each seasonal window. Predictions were generated on the logit (link) scale and back-transformed to the response scale to obtain the expected mean proportion surface. Wald 95% confidence intervals for the mean proportion were derived from the standard errors of the linear predictor. All statistical analyses described above were performed in R 4.5.1 (R Core Team, 2025).

### Extension of the gradient forest model to other localities

To test whether the Baltimore-Washington GF results generalized to regions sampled under different designs, we applied the same gradient forest approach — identical tree count, correlation threshold, and training/testing split — to Philadelphia and Chicago, modeling each locality independently, and additionally fit an aggregated model combining all three localities. As in Baltimore-Washington, these turnover models were fit to trap-night–level species proportion data and therefore included the weather predictors alongside the landscape predictors. We then compared the accumulated degree-day threshold at which compositional turnover was most rapid across the independent and aggregated models, to assess whether the Baltimore-Washington threshold recurred under different climates, landscapes, and *Culex* community compositions.

## Results

### Mosquito collection and identification

Across the BW study period, gravid traps collected 22,179 *Cx. pipiens*/*restuans* individuals (morphological identification) over 595 trap-nights, an average of 37.3 per trap-night. This is comparable to mean gravid-trap catches of these species reported from West Nile virus surveillance in the mid-Atlantic United States (32.3 and 64.0 per trap-night for species-resolved and total *Cx. pipiens*/*restuans* counts, respectively; Williams & Gingrich, 2007).

*Culex* specimens were successfully identified to species at a rate of 80%. A total of 9,789 mosquitoes were genotyped as *Cx. pipiens* or *Cx. restuans*. We genotyped 3,775 mosquitoes from Chicago (90.5% *Cx. pipiens*), 206 mosquitoes from Philadelphia (65.5% *Cx. pipiens*), and 5,808 from Baltimore-Washington (68.8% *Cx. pipiens*).

### Abundance modeling

Among single-predictor models of total *Culex* trap-night abundance in BW, elevation was the best-supported predictor (AIC = 5004.8; Table S2) and was positively associated with abundance in the single-predictor model (β = 0.34 ± 0.09 SE, *p* < 0.001). Adding percent impervious surface and its interaction with elevation improved model fit (AIC = 4999.2; Table S3). In this model, the interaction between elevation and impervious surface was strongly negative (β = −0.42 ± 0.12 SE, *p* < 0.001; Table 1), indicating that the positive association between elevation and abundance weakened and reversed as impervious surface increased.

Forward selection on the species-factor dataset identified percent impervious surface (AIC = 7796.7; Table S4) as the strongest single predictor and weekly mean temperature (AIC = 7709.0; Table S5) as the strongest second predictor. The impervious surface × temperature interaction (β = −0.07 ± 0.06 SE, *p* = 0.20) and the three-way species × impervious × temperature interaction (β = −0.09 ± 0.08 SE, *p* = 0.29) were non-significant and removed. When we added a directed species × elevation effect, model fit improved substantially (ΔAIC = 11.6; likelihood-ratio test χ² = 15.6, *p* < 0.001; Table S7), and we report this elevation-inclusive model as the species-factor model (Table 1). In this model, *Cx. pipiens* abundance increased with weekly mean temperature (β = 0.33 ± 0.07 SE, *p* < 0.001) while *Cx. restuans* declined (combined slope ≈ −0.47 ± 0.07 SE; species × temperature β = −0.80 ± 0.08 SE, *p* < 0.001; Fig. 2A). The two species diverged sharply along the urbanization gradient: *Cx. pipiens* abundance increased significantly with impervious surface (β = 0.36 ± 0.11 SE, *p* = 0.001), whereas *Cx. restuans* declined steeply (combined slope ≈ −0.95 ± 0.10 SE; species × impervious β = −1.31 ± 0.09 SE, *p* < 0.001; Fig. 2B). Both species increased with elevation, but *Cx. pipiens* did so more steeply (β = 0.48 ± 0.12 SE, *p* < 0.001) than *Cx. restuans* (combined slope ≈ 0.24 ± 0.12 SE; species × elevation β = −0.24 ± 0.08 SE, *p* = 0.005; Fig. 2C). The species-specific elevation slopes were essentially unchanged with vs without temperature in the model (Table S7b), indicating an elevation effect independent of temperature rather than a thermal proxy. Random effects in the species-factor model captured a moderate fraction of additional variance beyond the fixed effects (Table S1); inter-annual and within-season nightly heterogeneity were the largest sources of unmodeled variation, while between-site differences contributed less once environmental predictors were included. Site-level variation in the combined *Culex* model was substantially smaller than in the species-factor model (Table S1), indicating that the elevation × impervious surface interaction absorbed most of the persistent between-site structure in combined *Culex* abundance.

**Figure 2.**
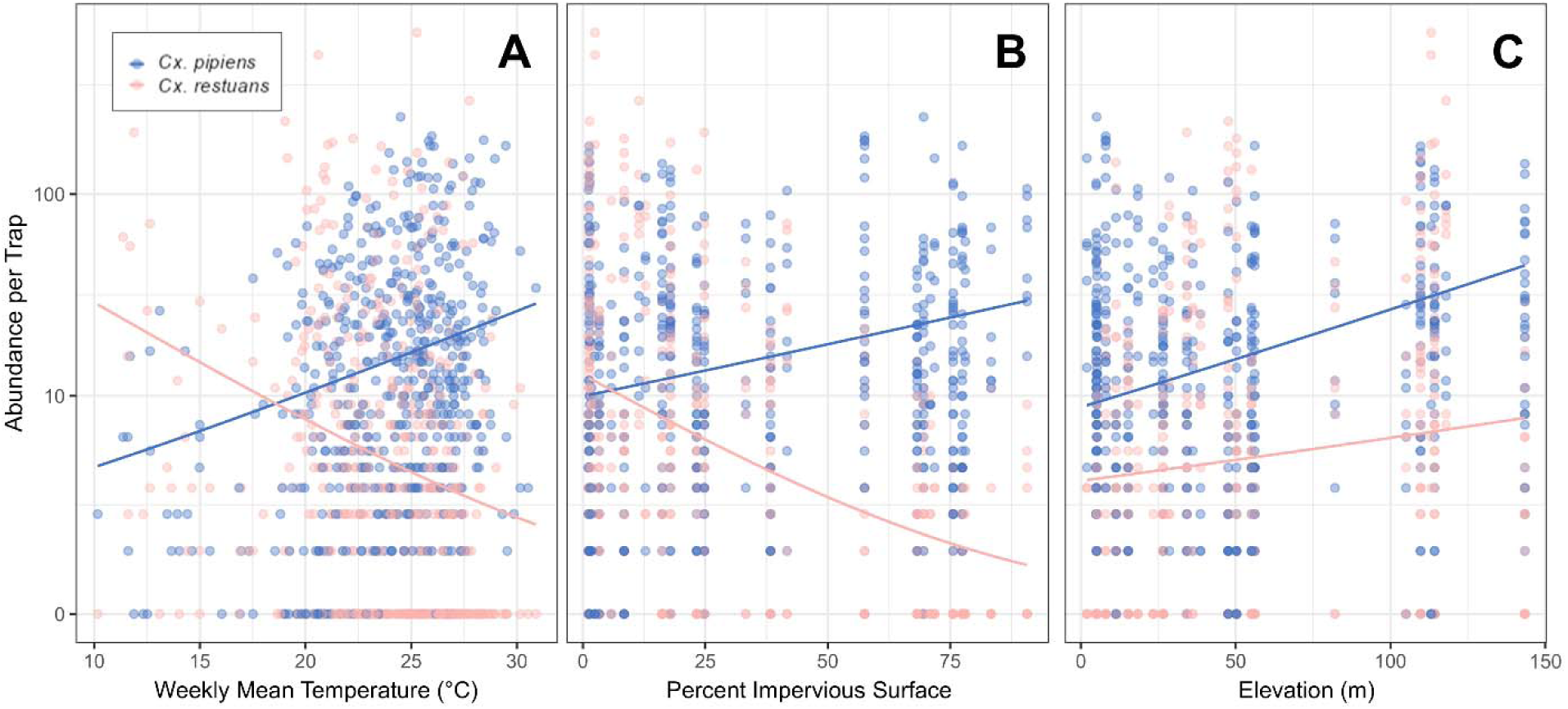
Observed trap-night abundance of *Cx. pipiens* and *Cx. restuans* across gradients of weekly mean temperature (A), percent impervious surface (B), and elevation (C) in Baltimore-Washington. Points represent individual trap-night counts. Lines show fitted relationships from the final negative binomial generalized linear mixed model, including fixed effects for species, weekly mean temperature, percent impervious surface, elevation, and the interactions between species and each environmental variable, with random intercepts for site, year, and trap night.

### Gradient forest modeling of species turnover in Baltimore-Washington

Accumulated degree-days was the second most important predictor of species turnover in the Baltimore-Washington GF model (superseded only by weekly mean temperature), with the transition from *Cx. restuans* to *Cx. pipiens* most rapid at 594 degree-days (21.4°C weekly mean temperature; Table 2). Among the landscape predictors, NDVI and percent impervious surface were the strongest drivers of turnover, consistent with the urbanization gradient identified in the abundance models above (Table 2).

**Table 2.** Overall model fit, R^2^-weighted predictor importance values, and values of most rapid turnover for the top eight most important predictors in each locality-specific GF model, plus the aggregated model. N indicates the number of trap-nights used as input for each model. BW includes data from 27 sites and six years (2019-2024), Philadelphia includes data from 65 sites and three years (2019-2021), and Chicago includes data from 30 sites and three years (2020-2022). ADD stands for accumulated degree-days.

| **BW (N = 576)**<br>R<sup>2</sup>: 0.36 |  |  | **Philadelphia (N = 71)**<br>R<sup>2</sup>: 0.26 |  |  | **Chicago (N = 403)**<br>R<sup>2</sup>: 0.39 |  |  | **All Locations (N = 1,052)**<br>R<sup>2</sup>: 0.43 |  |  |
| --- | --- | --- | --- | --- | --- | --- | --- | --- | --- | --- | --- |
| Predictor | Weighted importance | Most rapid turnover | Predictor | Weighted importance | Most rapid turnover | Predictor | Weighted importance | Most rapid turnover | Predictor | Weighted importance | Most rapid turnover |
| **Weekly Mean Temp** | 7.32 | 21.4°C | **Weekly Max Temp (Lagged)** | 0.98 | 30.1°C | **ADD** | 4.36 | 594 | **ADD** | 17.14 | 608 |
| **ADD** | 5.85 | 594 | **ADD** | 0.79 | 610 | **Day of Year** | 2.36 | 187 (July 6th) | **Day of Year** | 9.24 | 170 (June 19th) |
| **Day of Year** | 5.58 | 165 (June 14th) | **Weekly Mean Temp (Lagged)** | 0.74 | 21.3°C | **Weekly Max Temp (Lagged)** | 1.75 | 23.9°C | **Percent Impervious Surface** | 9.05 | 37.9% |
| **Weekly Max Temp** | 5.02 | 28.1°C | **Population Density** | 0.65 | 7.13 (1,250/km<sup>2</sup>) | **Weekly Max Temp** | 1.53 | 21.4°C | **Weekly Mean Temp** | 8.50 | 21.3°C |
| **NDVI** | 4.88 | 0.45 | **Housing Density** | 0.58 | 5.61 (273/km<sup>2</sup>) | **Weekly Rainfall (Lagged)** | 1.35 | 12.6mm | **Tree Canopy** | 8.21 | 17.4% |
| **Percent Impervious Surface** | 4.44 | 49.5% | **Weekly Mean Temp** | 0.54 | 22.3°C | **Weekly Mean Temp** | 1.23 | 21.9°C | **Weekly Max Temp** | 5.92 | 26.1°C |

### Spatial predictions in Baltimore-Washington

The BW-specific spatial GF models identified percent tree canopy, percent impervious surface, and NDVI as the three most important predictors of *Culex* relative abundance across space in both seasons (Table S9). The landscape values at which compositional turnover was most rapid - the maxima of the split-density-to-data-density ratio - were 19% tree canopy, 48% impervious surface, and an NDVI of 0.47 in the early season, and 51% tree canopy, 50% impervious surface, and an NDVI of 0.44 in the late season. The same three predictors contributed most strongly to PC1 in both seasons (Table S8), so PC1 can be interpreted as an urbanization-composite axis on a GF-derived biological-importance scale. PC1 explained 76.9% and 74.4% of the variance in the transformed predictor space in the early and late seasons, respectively (Table S8).

In both beta-binomial GLMs, *Cx. pipiens* relative abundance was strongly positively associated with PC1 (early season: β = 8.604 ± 1.638 SE, p < 0.001; late season: β = 9.188 ± 1.261 SE, p < 0.001). The large coefficient magnitudes reflect the GF-derived biological-importance scale of PC1, which lacks natural predictor units, rather than implausibly steep biological effects. Model uncertainty, quantified as the width of the 95% confidence interval, was greater in the early season—especially in highly urbanized areas (Fig. 3D). In the early season, the predicted proportion of *Cx. pipiens* was highest in downtown Washington and Baltimore and declined sharply with decreasing urbanization (Fig. 3A–C). From the early to the late season, predicted *Cx. pipiens* proportions increased across much of the region, particularly in suburban areas, where *Cx. pipiens* was predicted to exceed 50% of the community (Fig. 3E–G). Together, these analyses show that urbanization and seasonal temperature gradients jointly shape *Culex* community composition.

**Figure 3.**
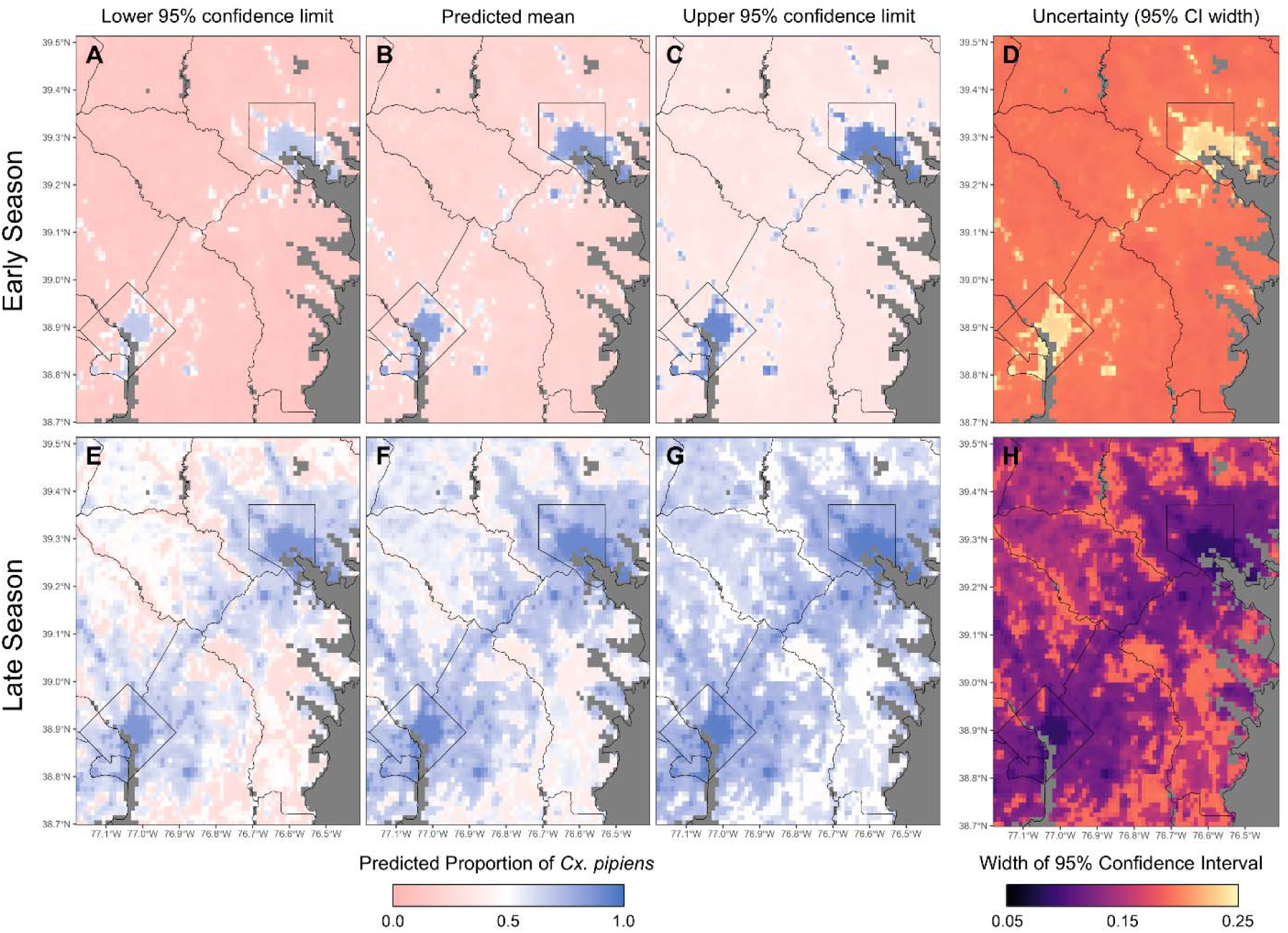
Spatial predictions of *Cx. pipiens* composition across the Baltimore-Washington region. Maps show the model-estimated proportion of *Cx. pipiens* among genotyped *Culex* individuals during the early (A–D) and late (E–H) seasonal windows. Panels display the lower 95% confidence limit (A, E), predicted mean proportion (B, F), and upper 95% confidence limit (C, G) derived from a beta-binomial generalized linear model relating species composition to PC1 (see Methods). Panels D and H show the width of the 95% confidence interval (upper − lower), representing uncertainty in the predicted mean proportion.

### Cross-regional transferability of the turnover threshold

The 594 degree-day threshold identified in BW was closely recovered in the other regions, each sampled independently and under different designs: turnover was most rapid at 594 degree-days in Chicago and 610 degree-days in Philadelphia, with the aggregated model yielding 608 degree-days (weekly mean temperature thresholds: Chicago 21.9°C, Philadelphia 22.3°C, aggregated 21.3°C; Table 2). That the same thermal threshold recurred across three regions with different climates, landscapes, and sampling designs is the strongest evidence in our data that the timing of the *Cx. restuans*-to-*Cx. pipiens* transition is governed primarily by temperature and is therefore transferable.

The landscape predictors that were influential in BW, by contrast, were not the same in the additional regions. Percent impervious surface, percent tree canopy, and vegetation index were among the least important predictors in both the Chicago and Philadelphia models, where weather predictors such as weekly maximum temperature and weekly rainfall were most strongly associated with turnover, though the aggregated model also identified percent impervious surface and percent tree canopy as important predictors (Table 2). Spatial turnover in Philadelphia was influenced by population and housing density, two additional measures of urban intensity. We computed pairwise correlations among all spatial predictors within each region (Tables S10–S12 for Baltimore-Washington, Chicago, and Philadelphia, respectively). Across Philadelphia sites, population density was moderately correlated with the land-cover urbanization predictors (|*r*| = 0.45–0.60 with percent impervious surface, percent tree canopy, and NDVI; Table S12), indicating that it partly indexes the urbanization gradient. Housing density was only weakly correlated with land cover (|*r*| ≤ 0.29), suggesting it captures a distinct metric of urban intensity. Although percent impervious surface and percent tree canopy ranked among the least important predictors in Philadelphia, the collinear population-density term was influential, so the urban signal there registered through demography rather than through land cover.

## Discussion

The climatic and landscape gradients that structure *Culex* vector communities are shifting, and the same pooled surveillance index that has served WNv programs for two decades is increasingly limited in what it can register about that change. In Baltimore-Washington, where balanced sampling supported species-specific abundance modeling, the two species diverged in opposite directions along both temperature and impervious-surface gradients, with urbanization shaping the spatial pattern of community composition. Across three eastern metropolitan regions, accumulated degree-days and weekly mean temperature drove a consistent seasonal turnover from *Cx. restuans* to *Cx. pipiens*, while urbanization shaped where that turnover was most pronounced wherever the sampling captured enough landscape variation to resolve it - through land cover in Baltimore-Washington and through population density in Philadelphia. These opposing species responses are the substrate from which species-resolved surveillance under environmental change can be built: they identify a transferable thermal threshold for the timing of community shifts, a spatial signature for where those shifts are most pronounced on the landscape, and a framework for anticipating how community composition may shift within these regions as local climate and land-use conditions change.

The relationship between accumulated degree-days, weekly mean temperature, and species composition was the strongest and most consistent signal across cities. The species-factor abundance model in Baltimore-Washington showed *Cx. pipiens* abundance rising with temperature while *Cx. restuans* declined (Table 1, Fig. 2A), and accumulated degree days was within the top two most important predictors of compositional turnover in all three regions (Table 2). The timing and duration of this temperature-driven seasonal transition have been associated with the timing and intensity of human West Nile virus cases (Lampman et al., 2006; Tokarz & Smith, 2020). Although this association does not by itself establish a causal pathway, it makes the transition a practical, climate-anchored target for anticipating the approaching period of elevated human risk. Turnover from *Cx. restuans* to *Cx. pipiens* was most rapid between 594 and 610 accumulated degree-days and at weekly mean temperatures of approximately 21–22 °C (Table 2), a relatively narrow thermal window over which the community shifts from the early-season *Cx. restuans* peak to the late-season *Cx. pipiens* peak. These patterns are consistent with extensive physiological evidence: *Cx. restuans* develops more rapidly at lower temperatures and exhibits higher mortality and reduced longevity above 25 °C, whereas *Cx. pipiens* immature stages develop more quickly and survive better at higher temperatures (Buth et al., 1990; Ciota et al., 2014; Madder et al., 1983).

Temperature interacted with landscape to shape spatial patterns of abundance in Baltimore-Washington. The two species diverged sharply along the urbanization gradient: *Cx. pipiens* abundance rose with increasing impervious surface, whereas *Cx. restuans* abundance declined steeply (Table 1, Fig. 2B). This opposing pattern is consistent with the broader literature: *Cx. pipiens* is well established as a generalist tolerant of urbanization (Arsenault-Benoit & Fritz, 2023; Bondo et al., 2023; Gorris et al., 2021; Johnson et al., 2015), while *Cx. restuans* is more strongly associated with vegetated habitats (Arsenault-Benoit & Fritz 2023; Gorris et al., 2021). The underlying mechanisms likely differ between species. *Cx. pipiens* exploits urban infrastructure - basements, sewers, and storm drains - for both oviposition during the breeding season and as overwintering hibernacula (Buffington, 1972; Hamer, 2014; Irwin 2008), and this use of subterranean habitats may buffer populations from extreme surface temperatures. *Cx. restuans*, in contrast, prefers more shaded oviposition sites associated with high tree canopy (Brust, 1990), which become scarce as impervious surface increases. The urban heat island effect may reinforce these differences, as impervious surfaces absorb and re-emit heat, elevating local temperatures relative to vegetated areas (Imhoff et al., 2010; Schwaab et al., 2021). Warmer urban conditions may favor the development and persistence of the more heat-tolerant *Cx. pipiens* over *Cx. restuans* (Ciota et al., 2014). Further, these ecological differences may reflect the species’ contrasting biogeographic histories. *Cx. restuans* is native to North America (Ross, 1964), whereas *Cx. pipiens* is generally regarded as introduced from the Old World (Fonseca et al., 2004; Harbach, 2012; Vinogradova, 2000). Native taxa are frequently filtered out by intensive development and anthropogenic disturbance, whereas non-native species that establish in human-dominated landscapes are, by definition, tolerant of such conditions (McKinney, 2006).

The species-factor model also showed that both *Culex* species increased in abundance with elevation, with *Cx. pipiens* responding more steeply than *Cx. restuans* (Table 1, Fig. 2C). This runs counter to declines in *Culex* abundance reported across some elevational gradients (Asigau et al., 2017; Eisen et al., 2008), though such relationships are not universal: in Hawaiian *Cx. quinquefasciatus*, abundance instead peaked at mid-elevation and was governed largely by the availability of larval habitat rather than by elevation per se (Villena et al., 2024). Those reported declines, moreover, were documented over broad montane gradients spanning hundreds to thousands of meters, across which elevation covaries strongly with temperature and whose upper reaches are cold enough to constrain mosquito development. Our gradient spans only tens of meters, and across our sites elevation and weekly mean temperature were only weakly correlated (r = −0.13), making the positive elevation effect largely independent of temperature. We therefore interpret elevation here not as a causal driver but as a proxy for unmeasured, physiographic habitat variation between the lower Coastal Plain and the higher Piedmont — an interpretation consistent with elevation ranking among the strongest predictors of *Cx. pipiens* occurrence at continental scales, on an axis distinct from temperature (Arora et al., 2022). Identifying the specific habitat features underlying this relationship will require finer-scale environmental data than the predictors used here.

The Philadelphia and Chicago datasets, drawn from existing operational surveillance and research collections rather than from designs optimized for parametric modeling of abundance, provided a test of whether the Baltimore-Washington thermal signal generalizes to systems sampled under different regimes. The three regions differ substantially in spatial extent, sampling intensity, and landscape context - Philadelphia collections emphasized broad spatial coverage over trap-night intensity within a compact urban footprint, while Chicago sampling covered a smaller geographic extent and yielded relatively few *Cx. restuans* - yet the thermal predictors driving compositional turnover converged across all three (Table 2). The specific landscape predictors did not transfer identically, but for different reasons in each region. In Chicago, no spatial predictor ranked among the six most important. Turnover was instead associated almost entirely with weather predictors, likely because the small sampling extent offered limited spatial and landscape variation relative to the weather variation in the dataset - the Chicago sites skewed heavily toward low tree canopy and high population density (Fig. S1). Consistent with these high-density, low-canopy conditions, Chicago collections were predominantly *Cx. pipiens* (90.5%), the species favored in heavily urbanized settings. In Philadelphia, the relevant covariate was instead population density, which is collinear with impervious surface and tree canopy (Table S12) and so indexes the same urbanization gradient through a different covariate. That the thermal threshold converged despite this much variation in sampling design and landscape context suggests it is a more general feature of the *Cx. pipiens*-*Cx. restuans* transition than any particular landscape predictor. The urbanization gradient itself is harder to evaluate: it shaped turnover in both Baltimore-Washington and Philadelphia - through land cover in one and population density in the other - whereas Chicago’s limited spatial variation may have reduced our power to detect significant changes due to the opportunistic sampling design. The gradient may therefore prove transferable as well, registering through whichever covariates best capture urbanization in a given region, but confirming this will require longer-term, spatially balanced sampling across a wider urbanization range.

### Implications for West Nile virus surveillance under environmental change

The opposing responses of *Cx. pipiens* and *Cx. restuans* along the temperature and urbanization gradients - both of which are themselves shifting under continuing climate change and urban expansion - underscore the need to understand the population dynamics of these two species and how those dynamics bear on surveillance and WNv risk prediction. The pooled “*Cx. pipiens/restuans*” abundance reported by most programs (Burgunder et al., 2026; Ebel et al., 2005; Harrington & Poulson, 2008; Johnson et al., 2015) sums two signals moving in opposite directions along each gradient: identical pooled totals can therefore correspond to very different species compositions, and to very different WNv transmission profiles, depending on where on the landscape and the seasonal calendar sampling occurs. Because human WNv transmission tracks the late-season *Cx. pipiens* phase most closely (Lampman et al., 2006; Tokarz & Smith, 2020), abundance models trained on the pooled index will under-resolve the component most directly relevant to risk. The species-specific environmental responses estimated here provide a framework in which existing pooled surveillance can be projected onto a species-resolved one, with the species-by-environment relationships used to translate routine pooled trap catches into species-specific abundance estimates anchored to local conditions.

The relatively narrow degree-day window (594-610 accumulated degree-days) over which compositional turnover is fastest provides an operational threshold that can serve as an anticipatory surveillance trigger rather than a retrospective description. Because the threshold can be evaluated from temperature data alone, vector-control agencies can forecast the impending growth of *Cx. pipiens* populations before they appear in traps, adjusting the timing and intensity of mosquito pool testing in advance rather than in reaction. A real-time degree-day forecast of this kind has been operating in central Illinois for two decades (Kunkel et al., 2006; Westcott et al., 2011), demonstrating that climate-based anticipatory surveillance is feasible to deploy. That the Baltimore-Washington threshold of 594 degree-days was closely matched in Chicago (594) and Philadelphia (610) - regions with different sampling designs, landscape contexts, and *Culex* community compositions - suggests the anticipatory use is likely to transfer across cities in the eastern United States, though local calibration would strengthen its application.

Spatially, the joint analysis of *Culex* composition identified the intermediate, roughly 20-50% impervious-surface landscapes characteristic of suburbs as exhibiting the largest seasonal shifts in community composition (Fig. 3). Concentrating added surveillance and early-season abatement in these landscapes - rather than spreading effort uniformly or focusing on the urban core, where composition shifts little - is likely to yield the largest returns for limited control resources, consistent with prior reports identifying suburban landscapes as recurrent hotspots of human West Nile virus cases (LaBeaud et al., 2008; Rochlin et al., 2011; Ruiz et al., 2007). Within-city risk stratification of this kind has been deployed effectively for other arboviral systems - for example, persistent dengue, chikungunya, and Zika hotspots have been identified across nine Mexican cities to prioritize *Aedes aegypti* control (Dzul-Manzanilla et al., 2021) - providing a methodological template for translating spatial heterogeneity in transmission into operational targeting decisions.

The value of these tools is increasing as the climatic and landscape gradients that structure the vector community themselves shift. Warming temperatures will move the 594–610 degree-day threshold earlier in the season, extending the *Cx. pipiens*-rich portion of the transmission season - and, because *Cx. pipiens* abundance rises with temperature while *Cx. restuans* declines, raising the share of late-season *Culex* catches composed of the species most strongly tied to human WNv transmission. Climate-driven northward expansion of *Culex* vectors has already been documented in Europe (Ainsworth, 2023; Erazo et al., 2024) and is projected for North America - including for both *Cx. pipiens* and *Cx. restuans* under multiple emissions scenarios (Gorris et al., 2021, Gorris et al., 2024; Samy et al., 2016), and continuing urban and suburban development is expected to expand, creating more of the preferred habitat of *Cx. pipiens* (Güneralp & Seto, 2013; Haba et al., 2025; Terando et al., 2014). Because the two species respond oppositely to both temperature and impervious surface, neither dimension of this change is recoverable from pooled abundance data: the same total catch can correspond to entirely different species compositions, and therefore to entirely different transmission profiles, depending on where on the gradients sampling occurs. Species-resolved environmental responses of the kind developed here are what adaptive surveillance under environmental change must be built on. Integrating species-specific abundance models with high-resolution projections of land-use change (Nowak & Greenfield, 2012) and climate (Gorris et al., 2021) will help anticipate how vector communities - and the WNv transmission risk they carry - will reorganize in the coming decades.

### Study limitations

Some limitations of our study warrant acknowledgment. Our molecular identification resolves *Cx. pipiens* from *Cx. restuans* but not bioforms within *Cx. pipiens* (form *pipiens*, form *molestus*, and hybrids), which differ in autogeny, host preference, and overwintering behavior (Mutebi & Savage, 2009). Resolving these bioforms would clarify the mechanisms underlying the urbanization signal we report, since the *molestus* and *pipiens* forms differ in their association with urban habitats. We nonetheless worked at the species level because it matches the scale at which routine surveillance operates and at which the tools we propose would be deployed. The 800 m resolution of the PRISM climate data masks fine-scale microclimatic heterogeneity that is increasingly recognized as an important driver of mosquito population dynamics in heterogeneous urban landscapes (Murdock et al., 2017; Sauer et al., 2021). Incorporating site-level or catch-basin temperature measurements would better resolve how fine-scale thermal environments interact with landscape structure to shape *Cx. pipiens* and *Cx. restuans* abundance. Additionally, approximately 20% of specimens could not be genotyped, primarily owing to incorrect morphological identification prior to extraction or poor DNA extraction quality. This is unlikely to have biased species abundance metrics, because the primer set distinguishes the two species reliably (Crabtree et al., 1995) and PCR failure under the rapid extraction protocol used here is not biased toward either species (Lévy et al., 2013; Rochlin et al., 2007). Finally, the species-environment relationships estimated here describe how *Cx. pipiens* and *Cx. restuans* respond to current temperature and urbanization gradients. Their use in anticipating community reorganization under future conditions assumes that those responses remain stationary as conditions move beyond the climatic and landscape range sampled in this study, a standard caveat for projection-based applications that local calibration and continued monitoring can help address.

## Conclusion

By integrating parametric (generalized linear mixed model) and non-parametric (Gradient Forest) approaches across multiple cities and years, this study shows that temperature governs the timing of seasonal turnover between two important West Nile virus vectors, that urbanization mediates the spatial pattern through opposing species responses, and that the resulting community structure can be distilled into operational tools for WNv surveillance: a transferable degree-day window for the timing of surveillance, a suburban target zone (roughly 20–50% impervious surface at a 1km resolution) for its spatial concentration, and species-resolved environmental responses that can be used to extract a species-specific signal from existing pooled surveillance and to anticipate how vector communities will reorganize under continuing climate and land-use change. Together they offer vector-control programs an anticipatory, spatially explicit framework for WNv surveillance - grounded in the ecology of the vector community itself and forward-looking under the environmental change reshaping it.

## Acknowledgments

This work was funded by NIH R01AI125622A to MLF. BG received partial support from CDC-TEC 6NU50CK000633-01-01. Maile Neel, Mark Clifton, and Linda Kothera also provided helpful comments that improved early drafts of this manuscript. We thank Christine Johnson, Linette Kingston, Helen Craig, Parker Grebe, Isabel Peñafiel, Ellen Osterman, and Nicole Guzman for help with mosquito collection and identification. Thank you to those who assisted us with site access: Al Greene (General Services Administration, retired), Arnaud Martin, Alex and Annie Kersey, Caroline Spiccioli, Rob McCann, David Serre, Josh Stift, Allison Dineen, Brenda Lee, Jeannine Dorothy (Maryland Department of Agriculture, retired), Patuxent Research Refuge (USFWS), Smithsonian Research and Education Center (SERC), the Smithsonian Institution, and University of Maryland Research and Education Centers at Beltsville, Upper Marlboro, and Clarksville. We also thank Brian Prendergast, who provided valuable insights into vector management practices in the State of Maryland. Artificial intelligence (GPT-4, OpenAI; Claude Opus 4.7, Anthropic) (used June 2025 and June 2026, respectively) was used to assist with R code for extracting cumulative importance values from the Gradient Forest turnover curves (see Methods) and for the species-factor model fit diagnostics reported in Table S1 (see Methods). All AI-assisted code was reviewed, tested, and validated by the authors, who are responsible for all content.

## Author Contributions

Benjamin P. Gregory, Arielle Arsenault-Benoit, and Megan L. Fritz conceived the ideas and designed methodology; Benjamin P. Gregory, Arielle Arsenault-Benoit, Patrick Irwin, Keith J. Price, and Bryn J. Witmier collected the data; Matthew C. Fitzpatrick consulted on GF modeling methodology; Benjamin P. Gregory and Arielle Arsenault-Benoit analyzed the data; Benjamin P. Gregory and Arielle Arsenault-Benoit led the writing of the manuscript. All authors contributed critically to the drafts and gave final approval for publication.

