## Appendix S1 for "Climate and urbanization structure West Nile virus vector communities: Tools for surveillance under environmental change"

**Authors:** *Benjamin P. Gregory, Arielle Arsenault-Benoit, Patrick Irwin, Keith J. Price, Bryn J. Witmier, Matthew C. Fitzpatrick, Megan L. Fritz*

**Supplemental Methods**

*Molecular Mosquito Identification*

Climate and urbanization structure West Nile virus vector communities: tools for surveillance under environmental change No. 69506, Qiagen Inc., Valencia CA, USA or Zymo Quick- DNA Miniprep Plus kit Cat #11-397B, Zymo Research, Irvine, CA) from the abdomen (excluding female spermathecae) in 2019 and 2020; or a rapid macerating procedure from the abdomen from 2021 through 2024 (Gloor & Engels 1992). Genomic DNA from Chicago and Philadelphia were all isolated using the rapid macerating procedure. Extracted DNA was amplified with a multiplexed polymerase chain reaction (PCR) targeting the 28S ribosomal subunit to identify *Culex pipiens s.l.* and *Culex restuans* as described by Crabtree et al. (1995) and Rochlin et al. (2007). This PCR also identifies *Culex salinarius*, but because gravid traps are biased against this species (Williams & Gingrich, 2007), we likely did not collect a representative sample of *Cx. salinarius* across the landscape. Therefore, any detected *Cx. salinarius* were excluded from analysis (N = 34).

PCRs were conducted according to the previously described protocols (Crabtree et al., 1995; Smith & Fonseca 2004). Two negative controls, a PCR negative in which purified water replaced genomic DNA, and an extraction negative, where the DNA extraction process was conducted without mosquito tissue, were always PCR amplified and electrophoresed alongside mosquito samples. Thermocycler conditions followed Crabtree et al. (1995) using a Bio-Rad T100 Thermal Cycler (Bio-Rad Life Sciences, Hercules, CA). Amplicons and controls were visualized on an agarose gel (2% w/v) alongside a 1Kb ladder (GeneRuler 1Kb Plus, ThermoScientific, Waltham MA) following gel electrophoresis at 120 v for 60 minutes, and identifications were made according to taxon-specific amplicon sizes.

*Justification for the directed species × elevation addition*

The species-specific abundance analysis presented in the main text combines two distinct model-building strategies: a two-stage forward selection by AIC that screens a broad set of candidate predictors, followed by a single directed addition of a species × elevation term to the AIC-selected model. The two stages address different inferential questions, and the directed addition was not pre-specified as a third forward-selection screen for the reasons set out here.

Forward selection is well suited to identifying the dominant axes of an unknown predictor space when the candidate set is large and the goal is parsimony — what subset of the available predictors best explains the response. For both the overall and species-specific datasets, forward selection identified compact two-predictor models that capture the strongest signals in the candidate set (Tables S2–S5): elevation interacted with percent impervious surface for overall *Culex* abundance, and species interacted with both percent impervious surface and weekly mean temperature for the species-split data. These selected models are appropriate end points of an exploratory screening procedure on the full candidate set.

Whether the two species diverge specifically along the elevational gradient, however, is a directed question with prior support rather than an open exploratory one. Elevation was the single most strongly supported predictor of overall *Culex* abundance (Table S2; AIC = 5004.8, β = 0.34, *p* < 0.001), establishing that the gradient carries appreciable abundance signal in this region. The forward-selected species model captures landscape (impervious surface) and climatic (temperature) axes but does not test whether the species responses to elevation differ from one another — a question on which the literature is ambiguous and which has direct bearing on the spatial structure of the vector community. The directed addition of a species × elevation term answers this specific question and reports the answer alongside the forward-selected effects.

We treated the directed addition as confirmatory rather than exploratory in three ways. First, we restricted the candidate to a single, named term (the species × elevation main effect plus interaction), rather than running a third forward-selection stage over the remaining predictor set, which would have inflated the model space and reintroduced the multiple-comparison problem that the two-stage procedure is designed to constrain. Second, we evaluated the term against three pre-specified criteria before retention: (i) it should not be collinear with the predictors already in the model; (ii) it should improve fit by a formal likelihood-ratio test; and (iii) its slopes should be stable to the inclusion or exclusion of weekly mean temperature, so that any observed effect is not a thermal proxy. Third, we fit the comparison set on a common complete-case subset of the data (n = 1,124 trap-night × species observations) so that the AIC and likelihood-ratio comparisons are valid.

All three criteria were met. Pairwise correlations among elevation, percent impervious surface, and weekly mean temperature were all below the conventional 0.7 collinearity threshold (strongest: elevation–impervious *r* = −0.50), and variance inflation factors were all below 1.4 (Table S6), well within the range supporting independent estimation of the three effects. Adding species × elevation to the AIC-best species model improved fit substantially (ΔAIC = 11.6; likelihood-ratio test χ²₂ = 15.55, *p* < 0.001; Table S7a). The per-species elevation slopes were essentially unchanged whether temperature was included or excluded from the model (*Cx. pipiens*: 0.484 with temperature vs 0.455 without; *Cx. restuans*: 0.245 vs 0.236; Table S7b), confirming that the elevation effect is not a temperature proxy. The elevation-inclusive model is reported as the focal species model (Table 2 of the main text).

This procedure separates the two inferential roles that the species model is asked to play. The forward-selected component identifies the dominant predictor pair from the full candidate set without a priori commitments, providing an unbiased screen of the available environmental information. The directed addition tests a single specific hypothesis — species-specific divergence along the dominant axis of overall abundance — against pre-specified criteria. Combining the two in a single procedure would have either inflated the screening multiplicity or required treating elevation as a priori (which it was not: its position in the analysis follows from the overall-abundance result, not from independent prior commitment). The procedure adopted here is the minimal one that answers both questions transparently.

*Predictor correlation structure and sampling-effort distributions*

To characterize the correlation structure among the spatial predictors used in the gradient forest models, we computed pairwise Pearson correlations among site-level predictor values (one observation per site) separately within each region (Baltimore–Washington, *n* = 27 sites; Chicago, *n* = 30; Philadelphia, *n* = 65; Tables S10–S12). These site-level correlations characterize the landscape sampled in each region and are distinct from the model-collinearity diagnostics in Table S6, which are computed on the trap-night × species modeling dataset to assess collinearity among the abundance-model predictors. To illustrate the range and evenness of sampling effort across predictors and regions (Figure S1), we plotted the distribution of each predictor across all trap-night collections rather than across unique sites, so that more heavily sampled sites contribute proportionally more to each distribution; this trap-night weighting reflects the realized allocation of sampling effort and shows, for example, the restricted range of the landscape predictors in the Chicago collections.

**Supplemental Figures**

**Figure S1.**

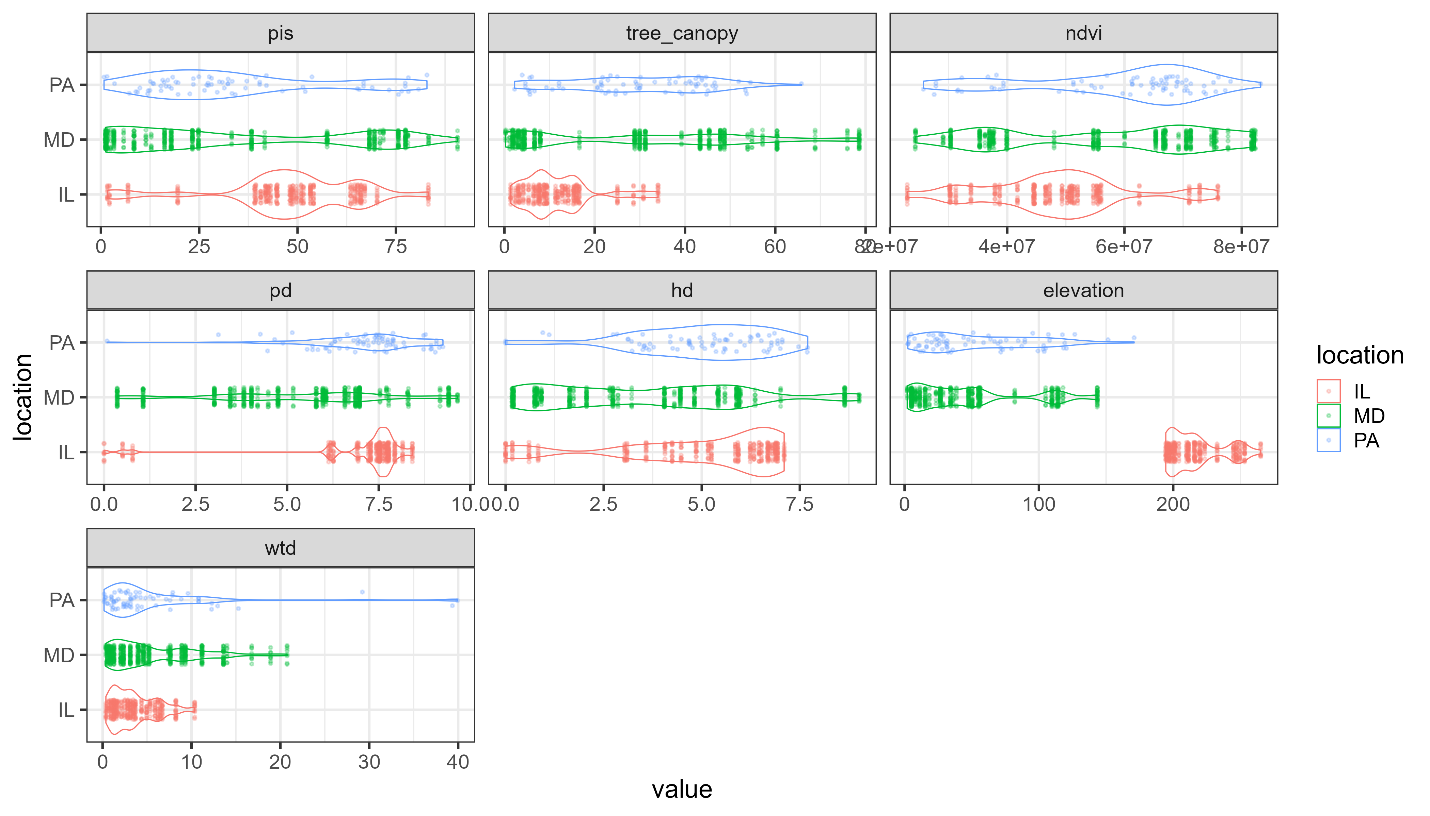

**Figure S1.** Distributions of landscape predictors across the three metropolitan regions (IL, Chicago; MD, Baltimore–Washington; PA, Philadelphia), shown as horizontal violins with overlaid points. Each point is a single trap-night, and the density curves are estimated across trap-nights rather than across unique sites, so sites and regions sampled more frequently contribute proportionally more to each curve; the distributions therefore reflect the realized distribution of sampling effort. Because the landscape predictors are constant within a site, repeated trap-nights at heavily sampled sites appear as points stacked at a single value. The x-axis is scaled independently within each panel. Panel abbreviations: pis, percent impervious surface; tree_canopy, percent tree canopy; ndvi, vegetation index; pd, population density; hd, housing density; elevation, elevation; wtd, water table depth.

**Supplemental Tables**

**Table S1.** Fit diagnostics for the two negative-binomial generalized linear mixed models (GLMMs) in Table 2 of the main text. Random-effect variance components are reported with corresponding standard deviations. Marginal *R*² (variance explained by fixed effects) and conditional *R*² (variance explained by fixed plus random effects) were computed using the trigamma approximation recommended for negative-binomial GLMMs (Nakagawa et al. 2017), implemented in the performance R package (Lüdecke et al. 2021). Both models produced mild non-convergence warnings under the bobyqa optimizer (max|grad| ≈ 0.03–0.05, tolerance = 0.002); refitting the focal species model with Nelder–Mead and nlminbwrap optimizers yielded identical fixed-effect estimates to four decimal places, indicating the warnings reflect tolerance thresholds rather than fit instability.

| **Quantity** | **Overall *Culex* model** | **Focal species model** |
| --- | --- | --- |
| N observations | 562 | 1,124 |
| N sites | 27 | 27 |
| N years | 6 | 6 |
| N trap-nights | 234 | 234 |
| Site variance σ² (SD) | 0.07 (0.26) | 0.17 (0.41) |
| Year variance σ² (SD) | 0.31 (0.56) | 0.34 (0.58) |
| Trap-night variance σ² (SD) | 0.37 (0.61) | 0.33 (0.58) |
| Negative binomial dispersion (θ) | 1.46 | 0.75 |
| Marginal *R*² (trigamma) | 0.09 | 0.27 |
| Conditional *R*² (trigamma) | 0.48 | 0.45 |

**Table S2.** Model selection results for single-predictor generalized linear mixed models (GLMMs) of overall *Culex* abundance in the Baltimore–Washington region. Models were ranked using Akaike’s Information Criterion (AIC); ΔAIC represents the difference in AIC relative to the lowest-AIC model within each model set (single- or two-predictor). Models with ΔAIC ≤ 2 are considered to have substantial support.

| Main Effect | AIC | ΔAIC |
| --- | --- | --- |
| Elevation | 5004.8 | 0.0 |
| Accumulated Degree Days | 5008.4 | 3.6 |
| Day of Year | 5008.7 | 3.9 |
| Weekly Rainfall (Lagged) | 5010.6 | 5.8 |
| Water Table Depth | 5013.2 | 8.3 |
| Percent Impervious Surface | 5014.2 | 9.4 |
| Weekly Rainfall | 5014.7 | 9.9 |
| Vegetation Index | 5015.2 | 10.4 |
| Housing Density | 5015.4 | 10.6 |
| Weekly Max Temperature | 5015.7 | 10.9 |
| Weekly Mean Temperature (Lagged) | 5015.8 | 10.9 |
| Population Density | 5015.8 | 11.0 |
| Percent Tree Canopy | 5015.9 | 11.1 |
| Weekly Max Temperature (Lagged) | 5016.0 | 11.2 |
| Weekly Mean Temperature | 5016.0 | 11.2 |

**Table S3.** Model selection results for two-predictor generalized linear mixed models (GLMMs) of overall Culex abundance in the Baltimore–Washington region. All models included fixed effects of elevation, the variable listed in the “Main Effect” column, and their interaction. Models were ranked using Akaike’s Information Criterion (AIC); ΔAIC represents the difference in AIC relative to the lowest-AIC model within each model set (single- or two-predictor). Models with ΔAIC ≤ 2 are considered to have substantial support.

| Main Effect | AIC | ΔAIC |
| --- | --- | --- |
| Percent Impervious Surface | 4999.2 | 0.0 |
| Day of Year | 4999.4 | 0.2 |
| Housing Density | 5000.1 | 0.9 |
| Accumulated Degree Days | 5001.0 | 1.7 |
| Water Table Depth | 5001.2 | 1.9 |
| Weekly Rainfall | 5002.9 | 3.7 |
| Weekly Rainfall (Lagged) | 5003.7 | 4.4 |
| Vegetation Index | 5005.2 | 6.0 |
| Percent Tree Canopy | 5005.9 | 6.6 |
| Weekly Max Temperature | 5007.2 | 8.0 |
| Weekly Max Temperature (Lagged) | 5007.7 | 8.4 |
| Population Density | 5008.0 | 8.8 |
| Weekly Mean Temperature (Lagged) | 5008.2 | 9.0 |
| Weekly Mean Temperature | 5008.3 | 9.1 |

**Table S4.** Model selection results for single-predictor species-specific generalized linear mixed models (GLMMs) of *Cx. pipiens* and *Cx. restuans* abundance in the Baltimore–Washington region. Models included species identity and its interaction with each environmental predictor. Models were ranked using Akaike’s Information Criterion (AIC); ΔAIC represents the difference in AIC relative to the lowest-AIC model within each model set.

| Main Effect | AIC | ΔAIC |
| --- | --- | --- |
| Percent Impervious Surface | 7796.7 | 0.0 |
| Vegetation Index | 7807.6 | 10.8 |
| Percent Tree Canopy | 7852.8 | 56.1 |
| Weekly Mean Temperature | 7894.7 | 97.9 |
| Weekly Mean Temperature (Lagged) | 7913.6 | 116.9 |
| Weekly Max Temp | 7922.3 | 125.6 |
| Weekly Max Temp (Lagged) | 7937.2 | 140.5 |
| Elevation | 7957.8 | 161.1 |
| Water Table Depth | 7958.3 | 161.5 |
| Accumulated Degree Days | 7961.1 | 164.3 |
| Day of Year | 7970.0 | 173.3 |
| Population Density | 7978.5 | 181.8 |
| Weekly Rainfall (Lagged) | 7987.5 | 190.7 |
| Weekly Rainfall | 7993.9 | 197.2 |
| Housing Density | 7994.1 | 197.3 |

**Table S5.** Model selection results for two-predictor species-specific generalized linear mixed models (GLMMs) of *Cx. pipiens* and *Cx. restuans* abundance in the Baltimore–Washington region. All models included fixed effects of species, percent impervious surface, the variable listed in the “Main Effect” column, and their interactions. Models were ranked using Akaike’s Information Criterion (AIC); ΔAIC represents the difference in AIC relative to the lowest-AIC model within each model set.

| Main Effect | AIC | ΔAIC |
| --- | --- | --- |
| Weekly Mean Temperature | 7709.0 | 0.0 |
| Weekly Max Temperature | 7718.8 | 9.8 |
| Weekly Mean Temperature (Lagged) | 7727.0 | 18.0 |
| Weekly Max Temperature (Lagged) | 7735.5 | 26.5 |
| Day of Year | 7779.3 | 70.4 |
| Percent Tree Canopy | 7783.2 | 74.2 |
| Accumulated Degree Days | 7783.8 | 74.8 |
| Elevation | 7786.5 | 77.6 |
| Vegetation Index | 7789.7 | 80.7 |
| Water Table Depth | 7794.5 | 85.5 |
| Weekly Rainfall (Lagged) | 7797.6 | 88.6 |
| Population Density | 7799.1 | 90.1 |
| Housing Density | 7799.3 | 90.4 |
| Weekly Rainfall | 7800.8 | 91.9 |

**Table S6.** Predictor collinearity diagnostics for the focal species model. (a) Pairwise Pearson correlations among the three focal predictors across the species-split modeling dataset (n = 1,124 complete-case trap-night × species observations). (b) Variance inflation factors (VIFs) for the three predictors, computed from a linear-regression analogue of the focal model fixed-effect structure. All pairwise |*r*| are below the 0.7 collinearity threshold and all VIFs are well below the conventional cutoff of 5, indicating that the three predictors carry largely independent information.

*(a) Pairwise Pearson correlations*

|  | **Elevation** | **Percent impervious surface** | **Weekly mean temperature** |
| --- | --- | --- | --- |
| Elevation | 1.000 | -0.499 | -0.133 |
| Percent impervious surface | -0.499 | 1.000 | 0.191 |
| Weekly mean temperature | -0.133 | 0.191 | 1.000 |

*(b) Variance inflation factors*

| **Predictor** | **VIF** |
| --- | --- |
| Elevation | 1.33 |
| Percent impervious surface | 1.36 |
| Weekly mean temperature | 1.04 |

**Table S7.** Justification of the directed species × elevation addition to the focal species model. (a) Likelihood-ratio comparison between the AIC-best two-predictor species model (impervious surface + weekly mean temperature, both interacted with species) and the focal model E2 (which adds species × elevation). (b) Per-species elevation slopes are essentially unchanged whether or not weekly mean temperature is included, confirming the elevation effect is independent of temperature rather than a thermal proxy. All models are negative-binomial GLMMs fit by maximum likelihood (Laplace approximation) with random intercepts for site, year, and trap-night, on the n = 1,124 complete-case observations of the species-split dataset.

*(a) Model comparison*

| **Model** | **Fixed-effect structure** | ***k*** | **AIC** | **ΔAIC** | **χ²** | **df** | ***p*** |
| --- | --- | --- | --- | --- | --- | --- | --- |
| Species model (no elevation) | species x (PIS + temperature) | 10 | 7711.0 | 11.6 | - | - | - |
| Focal model E2 | species x (elevation + PIS + temperature) | 12 | 7699.5 | 0.0 | 15.55 | 2 | <0.001 |

*k* = number of fixed-effect parameters. ΔAIC is computed relative to the focal model E2. χ², df, and *p* refer to the likelihood-ratio test comparing E2 to the no-elevation species model.

*(b) Stability of per-species elevation slopes with respect to temperature inclusion*

| **Species** | **E2 (temperature in):** **β ± SE** | **E1 (temperature out):** **β ± SE** |
| --- | --- | --- |
| *Cx. pipiens* | 0.484 ± 0.120 | 0.455 ± 0.125 |
| *Cx. restuans* | 0.245 ± 0.121 | 0.236 ± 0.126 |

Slopes are on the log link scale (a one-SD increase in scaled elevation multiplies expected abundance by exp(β)). The *Cx. restuans* slope is the sum of the elevation main effect and the species × elevation interaction; its SE is √(Var(β_main) + Var(β_int) + 2·Cov(β_main, β_int)).

**Table S8.** A summary of the Baltimore-Washington spatial-only principal component analyses for both the early and late seasons. The percentage of variance explained by each principal component (PC) (in parentheses) and the loadings for each GF-transformed predictor and each PC are shown.

| Predictor | Early Season | | | Late Season | | |
| --- | --- | --- | --- | --- | --- | --- |
|  | **PC1 (76.9%)** | **PC2 (10.4%)** | **PC3 (6.1%)** | **PC1 (74.4%)** | **PC2 (17.3%)** | **PC3 (3.3%)** |
| Percent Impervious Surface | 0.64 | -0.75 | -0.02 | -0.69 | -0.64 | 0.16 |
| NDVI | -0.61 | 0.52 | 0.52 | 0.38 | -0.02 | 0.01 |
| Percent Tree Canopy | -0.46 | -0.76 | -0.76 | 0.61 | -0.74 | 0.05 |
| Elevation | -0.05 | 0.14 | 0.39 | 0.04 | 0.22 | 0.61 |
| Water Table Depth | -0.05 | -0.01 | 0.01 | 0.07 | 0.02 | 0.77 |
| Population Density | 0.05 | -0.10 | -0.01 | -0.06 | -0.06 | 0.04 |
| Housing Density | 0.01 | -0.01 | -0.01 | -0.01 | -0.02 | 0.03 |

**Table S9.** Overall model fit and R^2^-weighted predictor importance values for the top six most important predictors in the Baltimore-Washington early and late season gradient forest models.

| **Early Season**  R^2^: 0.69 | | **Late Season**  R^2^: 0.62 | |
| --- | --- | --- | --- |
| Predictor | Weighted importance | Predictor | Weighted importance |
| **Percent Impervious Surface** | 0.57 | **Percent Tree Canopy** | 0.34 |
| **NDVI** | 0.52 | **Percent Impervious Surface** | 0.33 |
| **Percent Tree Canopy** | 0.46 | **NDVI** | 0.33 |
| **Elevation** | 0.23 | **Elevation** | 0.16 |
| **Water Table Depth** | 0.22 | **Population Density** | 0.10 |
| **Population Density** | 0.17 | **Water Table Depth** | 0.10 |
| **Housing Density** | 0.16 | **Housing Density** | 0.06 |

**Table S10.** Pairwise Pearson correlations among spatial predictors, Baltimore-Washington (n = 27 sites). Lower triangle shown; * p < 0.05, ** p < 0.01, *** p < 0.001.

|  | **Percent Impervious Surface** | **Percent Tree Canopy** | **NDVI** | **Population Density** | **Housing Density** | **Elevation** | **Water Table Depth** |
| --- | --- | --- | --- | --- | --- | --- | --- |
| **Percent Impervious Surface** | — |  |  |  |  |  |  |
| **Percent Tree Canopy** | -0.83*** | — |  |  |  |  |  |
| **NDVI** | -0.94*** | 0.95*** | — |  |  |  |  |
| **Population Density** | 0.45* | -0.30 | -0.38 | — |  |  |  |
| **Housing Density** | 0.12 | 0.00 | -0.03 | 0.81*** | — |  |  |
| **Elevation** | -0.56** | 0.45* | 0.55** | 0.05 | 0.29 | — |  |
| **Water Table Depth** | -0.34 | 0.39* | 0.41* | 0.15 | 0.27 | 0.15 | — |

**Table S11.** Pairwise Pearson correlations among spatial predictors, Chicago (n = 30 sites). Lower triangle shown; * p < 0.05, ** p < 0.01, *** p < 0.001.

|  | **Percent Impervious Surface** | **Percent Tree Canopy** | **NDVI** | **Population Density** | **Housing Density** | **Elevation** | **Water Table Depth** |
| --- | --- | --- | --- | --- | --- | --- | --- |
| **Percent Impervious Surface** | — |  |  |  |  |  |  |
| **Percent Tree Canopy** | -0.96*** | — |  |  |  |  |  |
| **NDVI** | -0.93*** | 0.93*** | — |  |  |  |  |
| **Population Density** | 0.39* | -0.36 | -0.19 | — |  |  |  |
| **Housing Density** | 0.26 | -0.32 | -0.06 | 0.77*** | — |  |  |
| **Elevation** | -0.37* | 0.21 | 0.26 | -0.34 | -0.28 | — |  |
| **Water Table Depth** | 0.02 | -0.10 | -0.05 | -0.04 | 0.08 | 0.59*** | — |

**Table S12.** Pairwise Pearson correlations among spatial predictors, Philadelphia (n = 65 sites). Lower triangle shown; * p < 0.05, ** p < 0.01, *** p < 0.001.

|  | **Percent Impervious Surface** | **Percent Tree Canopy** | **NDVI** | **Population Density** | **Housing Density** | **Elevation** | **Water Table Depth** |
| --- | --- | --- | --- | --- | --- | --- | --- |
| **Percent Impervious Surface** | — |  |  |  |  |  |  |
| **Percent Tree Canopy** | -0.86*** | — |  |  |  |  |  |
| **NDVI** | -0.95*** | 0.89*** | — |  |  |  |  |
| **Population Density** | 0.60*** | -0.45*** | -0.55*** | — |  |  |  |
| **Housing Density** | 0.29* | -0.20 | -0.26* | 0.74*** | — |  |  |
| **Elevation** | -0.40** | 0.27* | 0.42*** | -0.11 | 0.08 | — |  |
| **Water Table Depth** | -0.16 | 0.14 | 0.17 | -0.03 | -0.08 | 0.47*** | — |
